# SomaScan Bioinformatics: Normalization, Quality Control, and Assessment of Pre-Analytical Variation

**DOI:** 10.1101/2024.02.09.579724

**Authors:** Julián Candia

**Affiliations:** Translational Gerontology Branch, National Institute on Aging, National Institutes of Health, Baltimore, MD 21224, USA

## Abstract

SomaScan is an aptamer-based proteomics assay designed for the simultaneous measurement of thousands of human proteins with a broad range of endogenous concentrations. In its most current version released on November 1, 2023, the 11K SomaScan assay v5.0 is capable of measuring 10,776 human proteins covering major biological processes and disease areas, including cardiology, inflammation, neurology, and oncology. Here, I review bioinformatic approaches to perform normalization, quality control, and variability assessments.

## The SomaScan assay

SomaScan^1,2^ is a highly multiplexed, aptamer-based assay capable of simultaneously measuring thousands of human proteins broadly ranging from femto- to micro-molar concentrations. This platform relies upon a new generation of protein-capture SOMAmer (*Slow Offrate Modified Aptamer*) reagents^3^. SOMAmers are based on single-stranded, chemically-modified nucleic acids, selected via the so-called SELEX (*Systematic Evolution of Ligands by EXponential enrichment*) process, which is designed to optimize high affinity, slow off-rate, and high specificity to target proteins. These targets extensively cover major molecular functions including receptors, kinases, growth factors, and hormones, and span a diverse collection of secreted, intracellular, and extracellular proteins or domains. In recent years, SomaScan has increasingly been adopted as a powerful tool for biomarker discovery across a wide range of diseases and conditions, as well as to elucidate their biological underpinnings in proteomics and multi-omics studies^4–15^.

Concurrently with its wider adoption, SomaScan has expanded its proteome coverage by increasing the number of SOMAmers included in different versions of the assay, from roughly 800 SOMAmers in 2009, to 1,100 in 2012, 1,300 in 2015, 5,000 in 2018, 7,000 in 2020 and the most recent 11,000 protein assay available since late 2023 (Fig. 1). The first independent analysis of SomaScan normalization procedures and their variability was published by our teams at the U.S. National Institutes of Health (NIH) on the 1.1K and 1.3K assays^16^, later followed by technical reports from other laboratories^17–21^ and a recently updated assessment based on the 7K assay^22^.

**Figure 1.**
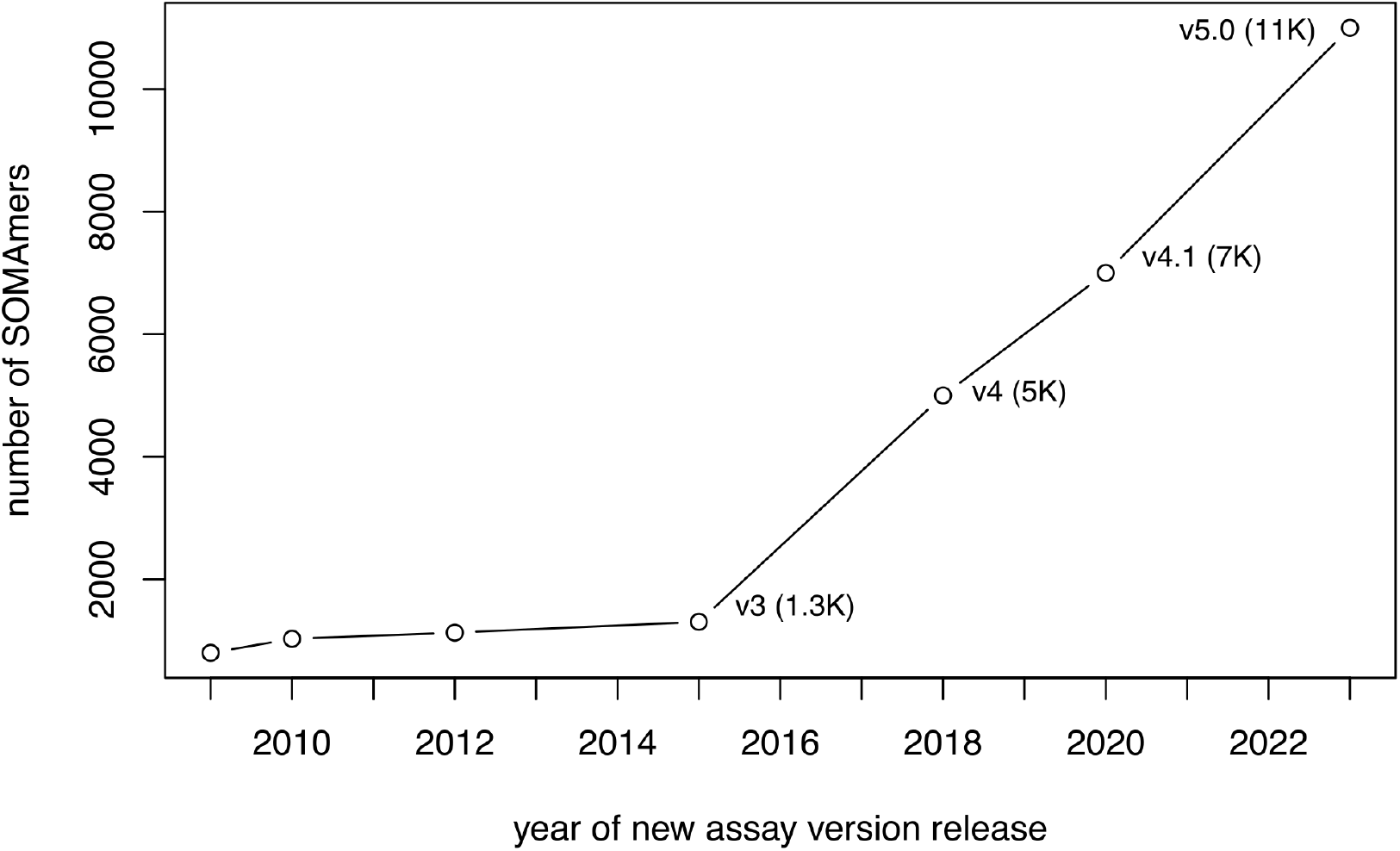
Timeline of SomaScan assay versions. Approximate timeline of assay version release dates showing the growth in the number of SOMAmers in each assay. Since 2015, the assay has been growing at a steady, approximately linear rate.

Table 1 shows the distribution of SOMAmer types in each SomaScan assay version. While most SOMAmers target human proteins, a small fraction of them targets mouse proteins. The remaining SOMAmers are different types of control, including twelve HCE (*Hybridization Control Elution*) SOMAmers used in the hybridization control normalization step (described below), as well as a few non-cleavable SOMAmers, non-biotin SOMAmers, spuriomers (designed as random, non-specific sequence motifs), and legacy SOMAmers targeting proteins from other species. In order to cover a broad range of endogenous concentrations, SOMAmers are binned into different dilution groups, namely 20% (1:5) dilution for proteins typically observed in the femto- to pico-molar range (which comprise about 80% of all human protein SOMAmers in the assay), 0.5% (1:200) dilution for proteins typically present in nano-molar concentrations (slightly below 20% of human protein SOMAmers in the assay), and 0.005% (1:20,000) dilution for proteins in micro-molar concentrations (about 2−3% of human protein SOMAmers in the assay). Although the proportions of SOMAmers across dilution groups have remained quite stable from version 5K onwards, it should be noticed that version 1.3K and earlier implemented a different scheme based on 40%, 1%, and 0.005% dilution groups. The human plasma or serum volume required is 55 *µ*L per sample.

**Table 1.** Distribution of SOMAmer types.

| SOMAmer Type vs Assay | 11K | 7K | 5K | 1.3K |
| --- | --- | --- | --- | --- |
| Human Protein | 10776 | 7289 | 4979 | 1302 |
| Mouse Protein | 233 | 233 | 228 | - |
| Hybridization Control Elution | 12 | 12 | 12 | 12 |
| Non-biotin | 10 | 10 | 10 | - |
| Non-cleavable | 4 | 4 | 4 | - |
| Spuriomer | 20 | 20 | 20 | - |
| Other | 28 | 28 | 31 | 8 |

SOMAmers are uniquely identified by their “SeqId”, but the relation between SOMAmers and annotated proteins is not one-to-one. In the 11K assay, for instance, the 10,776 SOMAmers that target human proteins are mapped to 9,609 unique UniProt IDs, 9,550 unique Entrez gene IDs, and 9,604 unique Entrez gene symbols. In the 7K assay, the 7,289 SOMAmers that target human proteins are mapped to 6,383 unique UniProt IDs, 6,378 unique Entrez gene IDs, and 6,383 unique Entrez gene symbols. Among cases where two or more SOMAmers share the same annotated target, the observed RFU correlation distribution is bi-modal, with peaks around *r* ≈ 0 and *r* ≈ 1^22^. While the *r* ≈1 peak simply indicates the expected redundancy of SOMAmers that bind to the same target, the *r* ≈ 0 peak may be due to SOMAmers that bind to different proteoforms annotated under the same protein target name, although they may also be due to artifacts such as cross-reactivity and non-specific binding. All of the SOMAmers in the 7K and 5K SomaScan assays, as well as most of those in the 1.3K version, are included in the newest 11K assay (Fig. 2). Data processing and delivery by SomaLogic is in the `adat` format, which is described in SomaLogic’s public GitHub repository (https://github.com/SomaLogic) and has remained consistent across assay versions. The data normalization procedures described below are equally applicable to all assay versions.

**Figure 2.**
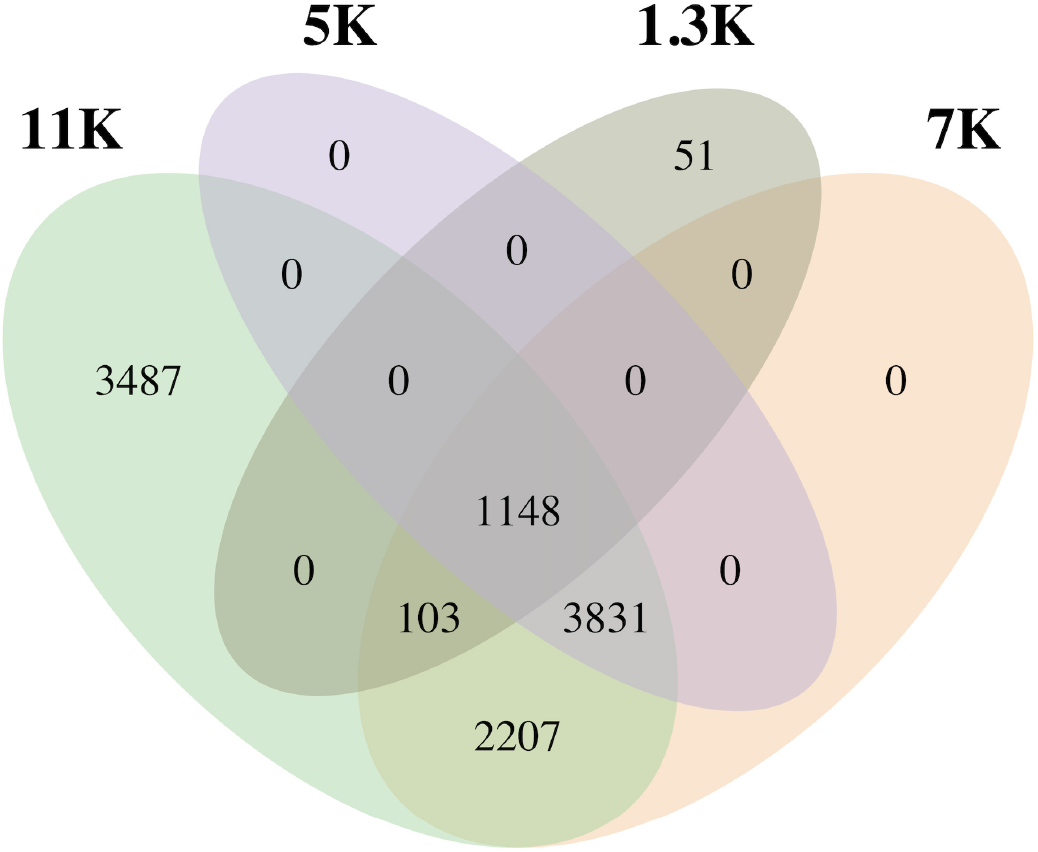
Human protein SOMAmers across SomaScan versions. Venn diagram showing the SOMAmer overlap (based on “SeqId” identifiers) between the 1.3K, 5K, 7K, and 11K SomaScan assays.

The experimental workflow, summarized in Fig. 3, consists of a sequence of steps, namely: (1) SOMAmers are synthesized with a fluorophore, photocleavable linker, and biotin; (2) diluted samples are incubated with dilution-specific SOMAmers bound to streptavidin beads; (3) unbound proteins are washed away, and bound proteins are tagged with biotin; (4) UV light breaks the photocleavable linker, releasing complexes back into solution; (5) non-specific complexes dissociate while specific complexes remain bound; (6) a polyanionic competitor is added to prevent rebinding of non-specific complexes; (7) biotinylated proteins (and bound SOMAmers) are captured on new streptavidin beads; and (8) after SOMAmers are released from the complexes by denaturing the proteins, fluorophores are measured following hybridization to complementary sequences on a microarray chip. The fluorescence intensity detected on the microarray, measured in RFU (*Relative Fluorescence Units*), is assumed to reflect the amount of available epitope in the original sample.

**Figure 3.**
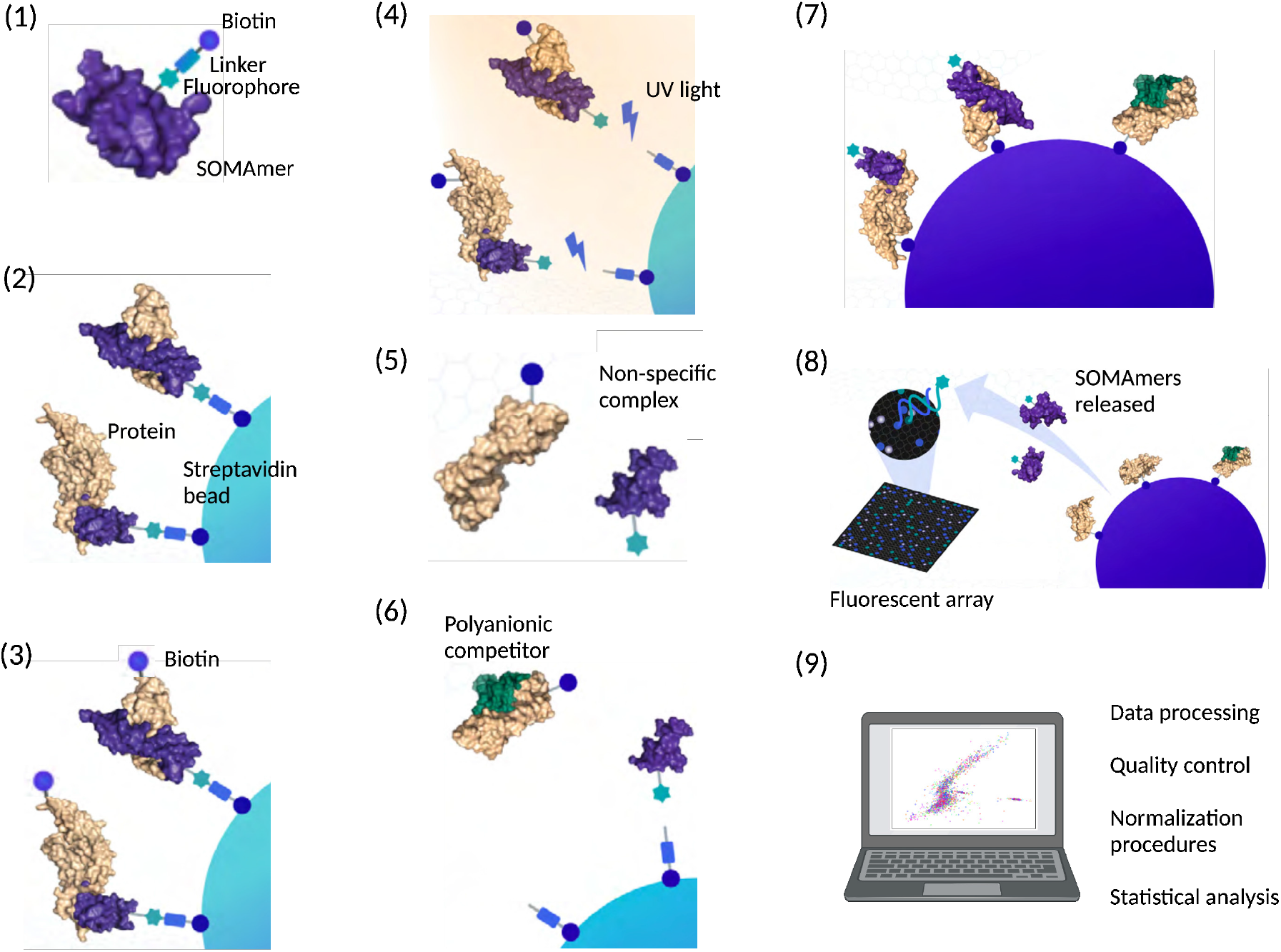
Workflow of the SomaScan assay. (1) SOMAmers are synthesized with a fluorophore, photocleavable linker, and biotin; (2) diluted samples are incubated with dilution-specific SOMAmers bound to streptavidin beads; (3) unbound proteins are washed away, and bound proteins are tagged with biotin; (4) UV light breaks the photocleavable linker, releasing complexes back into solution; (5) non-specific complexes dissociate while specific complexes remain bound; (6) a polyanionic competitor is added to prevent rebinding of non-specific complexes; (7) biotinylated proteins (and bound SOMAmers) are captured on new streptavidin beads; and (8) after SOMAmers are released from the complexes by denaturing the proteins, fluorophores are measured following hybridization to complementary sequences on a microarray chip. Upon completion of all experimental steps, (9) the bioinformatic analysis proceeds. Adapted from SomaLogic’s Technical Note SL00000572 and created with BioRender.

Samples are organized in 96*−*well plates, each plate consisting of buffer wells (reagent blanks with no sample material added), calibrator and QC samples (which are provided by SomaLogic from pooled healthy donor controls), and the experimental samples of interest. SomaScan users may be interested in adding their own set of control samples to the plate design, which would allow them to bridge across studies independently from SomaLogic-provided controls. On the one hand, SomaLogic’s controls may change over time, making it difficult to compare between studies run at different times. On the other hand, owning a set of control samples may allow users to use other omics technologies, including other proteomic assays such as mass spectrometry and Olink^23^, for further validation. As a standard practice, our labs at the NIH have implemented the use of a control sample derived from pooled healthy donors, which is run on every plate with 3 to 4 technical replicates per plate.

## Data Normalization Procedures

Raw data, as obtained after aggregation from slide-based hybridization microarrays, exhibit intra-plate nuisance variance due to differences in loading volume, leaks, washing conditions, etc, which is then compounded with batch effects across plates. In order to account for intra- and inter-plate variability of buffer, calibrator, QC, and experimental samples, here we consider a sequence of steps, whose main elements are hybridization normalization, median signal normalization, plate-scale normalization, and inter-plate calibration. Each of these steps generates scale factors at different levels: plate-specific, by SOMAmer dilution group, SOMAmer-specific, by sample type, and combinations thereof. Besides removing technical variability, these scale factors can be used as quality control flags at the plate-, sample-, and SOMAmer-levels^8^. A summary of these data normalization procedures, explained below in detail, is shown in Table 2. To keep a consistent mathematical notation in the expressions below, wells are indicated by Latin sub-indices and SOMAmers by Greek sub-indices. Plate index, sample type, and SOMAmer groupings are denoted by super-indices. We use *well* and *sample* interchangeably, although it should be noticed that no sample material is added to buffer wells; also, depending on context, *sample* may refer specifically to the experimental samples of interest, thus excluding control sample types such as buffer, calibrator, and QC.

**Table 2.** Summary of normalization steps.

| Step | Name | Abbreviation | Scale Factors |
| --- | --- | --- | --- |
| 0 | Raw | raw | none |
| 1 | Hybridization control normalization | hyb | well-specific |
| 2 | Median signal normalization on calibrators | hyb.msnCal | calibrator- and dilution-specific |
| 3 | Plate-scale normalization | hyb.msnCal.ps | plate-specific |
| 4 | Inter-plate calibration | hyb.msnCal.ps.cal | plate- and SOMAmer-specific |
| 5 | Median signal normalization on all sample types | hyb.msnCal.ps.cal.msnAll | well-specific (grouped by type) |

As we describe each normalization step, we also include R code chunks that implement each of those steps. We assume that the raw `adat` file is available to the user and that it has been split into three different plain-text files, namely: (i) “samples.txt”, with headers and one sample per row, providing metadata information on each well in the study; (ii) “somamers.txt”, with headers and one SOMAmer per row, providing metadata information on each SOMAmer in the assay; and (iii) “RFU.raw.txt”, without headers, samples as rows and SOMAmers as columns, providing the measured raw RFU values for each well and SOMAmer in the study. The ordering of samples and SOMAmers in “RFU.raw.txt” must be consistent with that of “samples.txt” and “somamers.txt”, respectively. Files in the `adat` format can be opened and manipulated programmatically via custom scripts, using the `SomaDataIO` R package available from SomaLogic’s public GitHub repository (https://github.com/SomaLogic), or using previously developed open-source tools^24,25^; however, it is often simple enough to open the `adat` file in Excel or similar software, then copy-paste the three separate regions corresponding to sample metadata, SOMAmer metadata (that needs to be transposed in order to display SOMAmers row-wise) and RFU matrix, and save them separately as tab-delimited plain-text files. Fig. 4 shows an example of the input data files required for this normalization pipeline.

**Figure 4.**
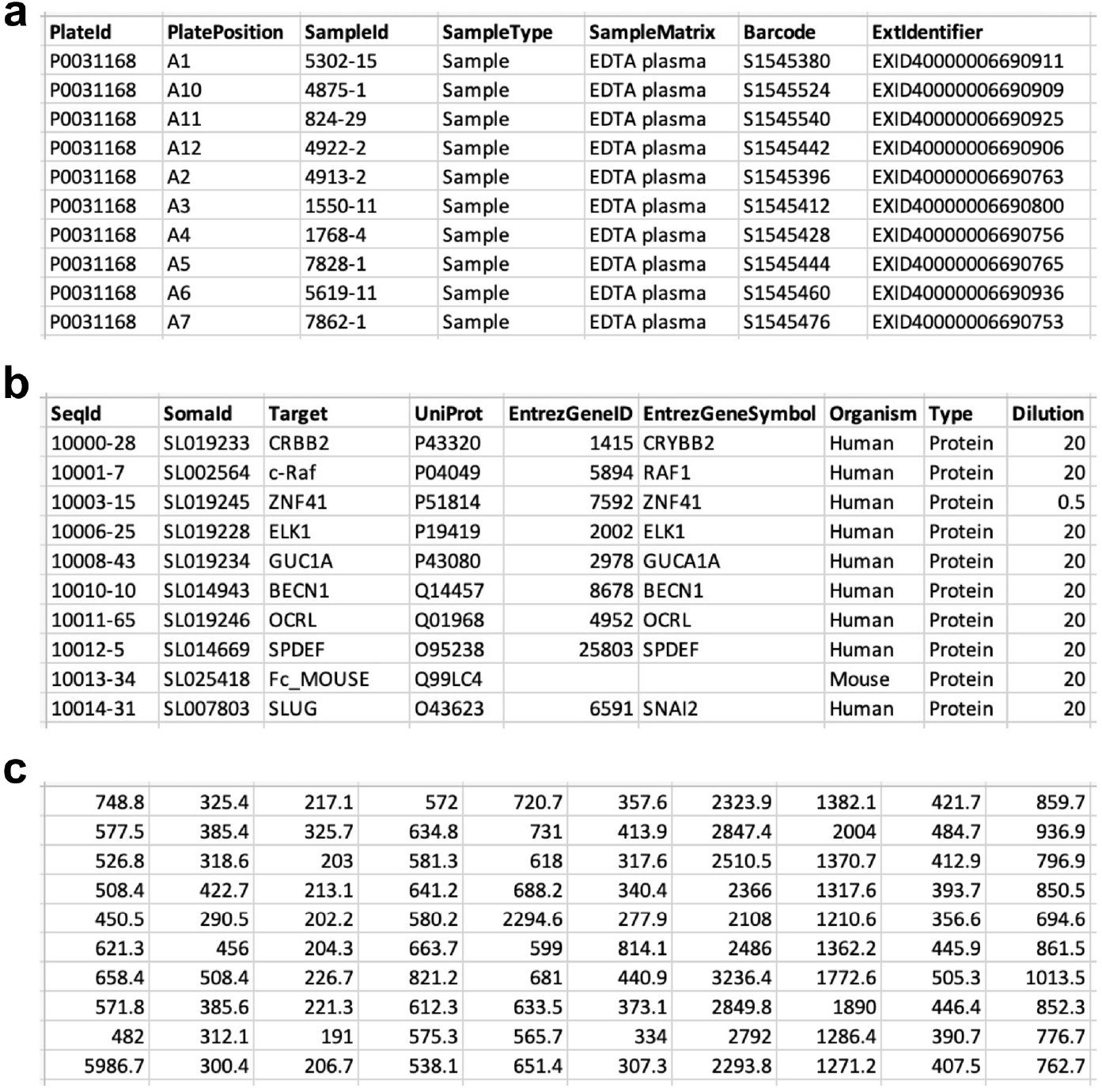
Input data files for the normalization pipeline. **a** “samples.txt”, with headers and one sample per row, provides metadata information on each well in the study (including buffer, QC, calibrator, and experimental samples). **b** “somamers.txt”, with headers and one SOMAmer per row, provides metadata information on each SOMAmer in the assay (including controls such as Hybridization Control Elution (HCE) SOMAmers). **c** “RFU.raw.txt”, without headers, samples as rows and SOMAmers as columns, provides the measured raw RFU values for each well and SOMAmer in the study (in the same order as the sample and SOMAmer metadata files).

The following code chunk reads the input datasets and sets up the variables needed to implement each of the normalization steps described below. Notice that, for the sake of clarity, we avoid using variable names that overlap with names of core R functions (e.g. `sample`). The code presented here was written with an emphasis on readability but not optimized for computing performance. Since `adat` files are much smaller than typical datasets generated by other omics and bioinformatic tasks, further optimization was not deemed necessary.

~~~
1 norm=**c**(“raw”,”hyb”,”hyb.msnCal”,”hyb.msnCal.ps”,”hyb.msnCal.ps.cal”,”hyb.msnCal.ps.cal.msnAll”)
2 n**_**norm = **length**(norm)
3 RFU = **vector**(“list”,n**_**norm)
4 sampl=**as.matrix**(**read.table**(“samples.txt”,header=T,**quote**=““,**comment.char**=““,sep=“\t”))
5 somamers=**as.matrix**(**read.table**(“somamers.txt”,header=T,**quote**=““,**comment.char**=““,sep=“\t”))
6 RFU[[1]]=**as.matrix**(**read.table**(“RFU.raw.txt”,header=F,sep=“\t”))
7 n**_**sampl = **nrow**(sampl)
8 n**_**somamer = **nrow**(somamer)
9 plate = **unique**(sampl[,”PlateId”])
10 n**_**plate = **length**(plate)
11 dil = **c**(0.005,0.5,20)
12 dil**_**lab = **c**(“0**_**005”,”0**_**5”,”20”)
13 n**_**dil = **length**(dil)
~~~

### 1. Hybridization control normalization (hyb )

Hybridization control normalization is designed to adjust for nuisance variance on the basis of individual wells. Each well contains *n*_*HCE*_ = 12 HCE (*Hybridization Control Elution*) SOMAmers at different concentrations spanning more than 3 orders of magnitude. By comparing each observed HCE probe to its corresponding reference value and then calculating the median over all HCE probes, we obtain the scale factor for the *i*-th well, i.e.

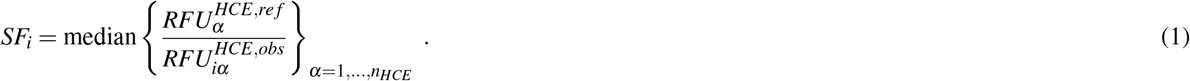

Notice that this normalization step is performed independently for each well; once the scale factor is determined, all SOMAmer RFUs in the well are multiplied by the same scale factor. Instead of using an external reference, a plate-specific internal reference can be determined by the median across the *n*_*s*_ = 96 wells in the plate, i.e.

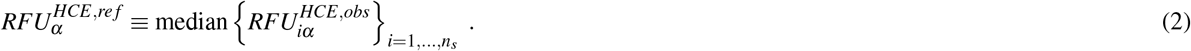

The following code chunk implements the hybridization control normalization step:

~~~
14 sel**_**HCE = somamer[,”Type”]==“Hybridization Control Elution”
15 n**_**HCE = **sum**(sel**_**HCE)
16 RFU**_**HCE = RFU[[1]][,sel**_**HCE]
17 RFU[[2]] = RFU[[1]]
18 SF**_**hyb = **rep**(NA,n**_**sampl)
19 HCE**_**ratio = **matrix**(**rep**(NA,n**_**sampl^**\***^n**_**HCE),**ncol**=n**_**HCE)
20 **for** (i**_**plate in 1:n**_**plate) {
21     sampl**_**sel = sampl[,”PlateId”]==plate[i**_**plate]
22     hyb**_**intra**_**ref = **apply**(RFU**_**HCE[sampl**_**sel,],2,**median**)
23     HCE**_**ratio**_**plate = HCE**_**ratio[sampl**_**sel,]
24     SF**_**hyb**_**plate = SF**_**hyb[sampl**_**sel]
25     RFU**_**plate = RFU[[2]][sampl**_**sel,]
26     **for** (i**_**sampl in 1:**sum**(sampl**_**sel)) {
27         HCE**_**ratio**_**plate[i**_**sampl,] = hyb**_**intra**_**ref**/**RFU**_**plate[i**_**sampl,sel**_**HCE]
28         SF**_**hyb**_**plate[i**_**sampl] = **median**(HCE**_**ratio**_**plate[i**_**sampl,])
29         RFU**_**plate[i**_**sampl,] = SF**_**hyb**_**plate[i**_**sampl]^**\***^RFU**_**plate[i**_**sampl,]
30     }
31     HCE**_**ratio[sampl**_**sel,] = HCE**_**ratio**_**plate
32     SF**_**hyb[sampl**_**sel] = SF**_**hyb**_**plate
33     RFU[[2]][sampl**_**sel,] = RFU**_**plate
34 }
~~~

### 2. Median signal normalization on calibrators (hyb.msnCal)

Median signal normalization is an intra-plate normalization procedure performed within wells of the same sample class (i.e. separately for buffer, QC, calibrator, and experimental samples) and within SOMAmers of the same dilution group. It is intended to remove sample-to-sample differences in total RFU brightness that may be due to differences in overall protein concentration, pipetting variation, variation in reagent concentrations, assay timing, and other sources of variability within a group of otherwise comparable samples. Since RFU brightness differs significantly across SOMAmers, median signal normalization proceeds in two steps. First, the median RFU of each SOMAmer is determined (across all samples of the same sample type) and sample RFUs are divided by it. The ratio corresponding to the *i*-th sample and *α*-th SOMAmer is thus given by

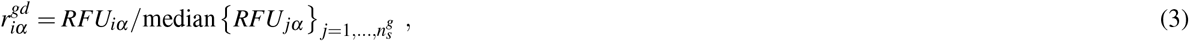

where indices *g* and *d* denote sample type and SOMAmer dilution groupings, respectively. Then, the scale factor associated with the *i*-th sample is determined as the inverse of the median ratio for that sample across all SOMAmers in the dilution group:

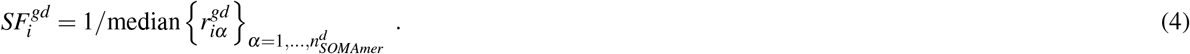

To median-normalize the *i*-th sample, then, all its SOMAmer RFUs in the same dilution group are multiplied by this scale factor. This procedure is illustrated by Fig. 5. As discussed in our previous work^16^, performing median signal normalization on experimental samples *before* inter-plate calibration presents the risk of enhancing plate-to-plate differences. Thus, in this step, we restrict median signal normalization to calibrators only.

**Figure 5.**
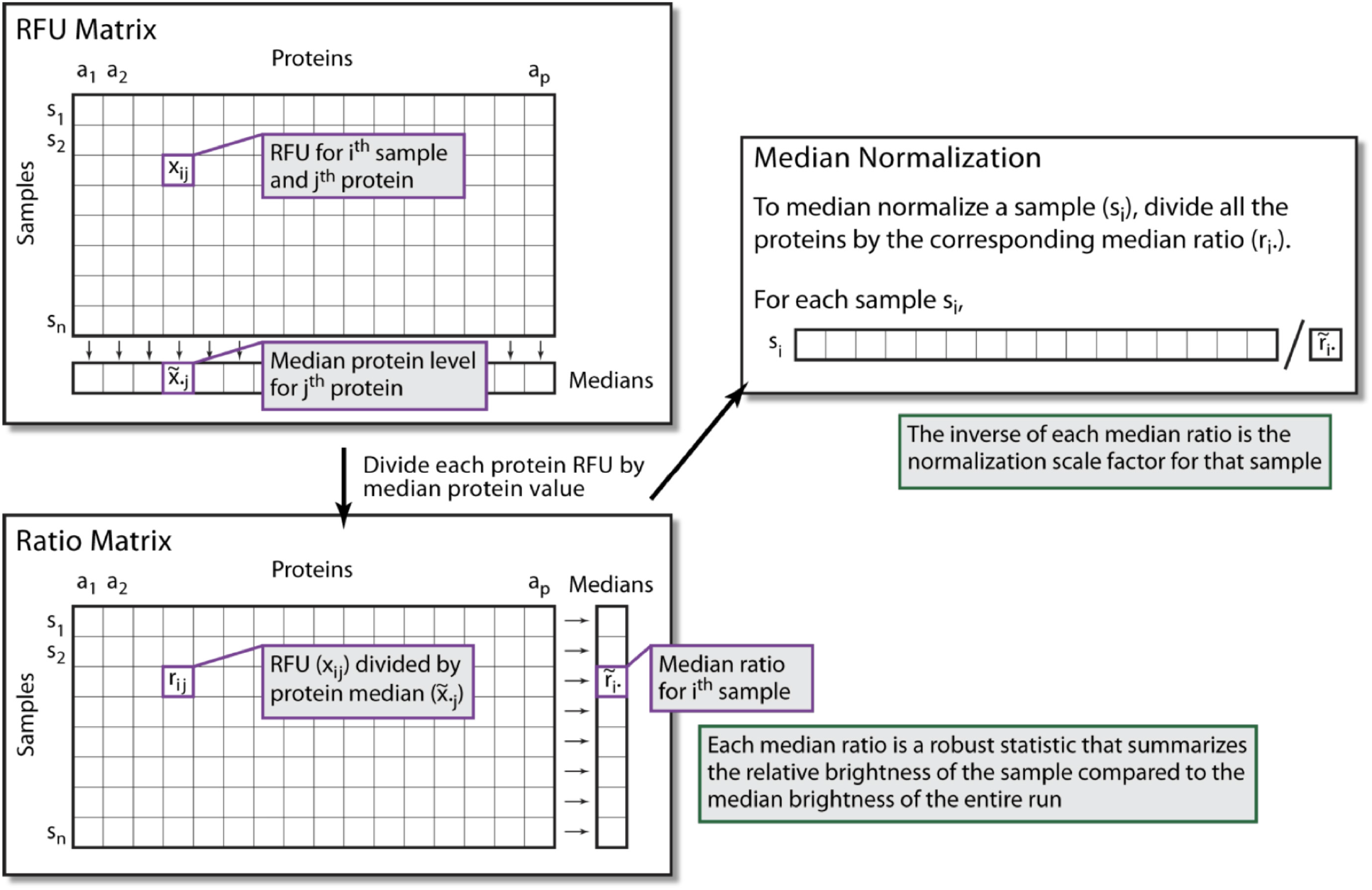
Median signal normalization. An RFU sub-matrix is defined by all samples of the same class (i.e. separately for buffer, QC, calibrator, and experimental samples) and all SOMAmers within the same dilution group. For each SOMAmer, the median RFU level is calculated across all samples (top left). By dividing each RFU measurement by the corresponding SOMAmer median, we obtain the ratio matrix; then, for each sample, a median ratio is calculated (bottom left). The median normalization scale factor associated to each sample is the inverse of the median ratio. To median-normalize each sample, all SOMAmers within the target dilution group are multiplied by its corresponding scale factor. Adapted from image courtesy of Darryl Perry (SomaLogic).

The following code chunk implements the median signal normalization step on calibrators:

~~~
35 RFU[[3]] = RFU[[2]]
36 SF**_**medNormInt = **matrix**(**rep**(1,n**_**sampl^**\***^n**_**dil),**ncol**=n**_**dil)
37 sampl**_**type = **c**(“Calibrator”)
38 n**_**sampl**_**type = **length**(sampl**_**type)
39 **for** (i**_**plate in 1:n**_**plate) {
40      sel1 = sampl[,”PlateId”]==plate[i**_**plate]
41      **for** (i**_**sampl**_**type in 1:n**_**sampl**_**type) {
42           sel2 = sampl[,”SampleType”]==sampl**_**type[i**_**sampl**_**type]
43           indx**_**sampl = **which**(sel1**&**sel2)
44           **for** (i**_**dil in 1:n**_**dil) {
45               sel**_**somamer = **as.numeric**(somamer[,”Dilution”])==dil[i**_**dil]
46               dat = RFU[[3]][indx**_**sampl,sel**_**somamer,**drop**=F]
47               dat2 = dat
48               **for** (i in 1:**ncol**(dat2)) {
49                    dat2[,i] = dat2[,i]**/median**(dat2[,i])
50               }
51               **for** (i in 1:**nrow**(dat)) {
52                    SF**_**medNormInt[indx**_**sampl[i],i**_**dil] = 1**/median**(dat2[i,])
53                    dat[i,] = dat[i,]^**\***^SF**_**medNormInt[indx**_**sampl[i],i**_**dil]
54               }
55               RFU[[3]][indx**_**sampl,sel**_**somamer] = dat
56           }
57      }
58 }
~~~

### 3. Plate-scale normalization (hyb.msnCal.ps)

Plate-scale normalization aims to control for variance in total signal intensity from plate to plate. No protein spikes are added to the calibrator; the procedure solely relies on the endogenous levels of each protein within the set of calibrator replicates.

For the *α*-th SOMAmer on the *p*-th plate,

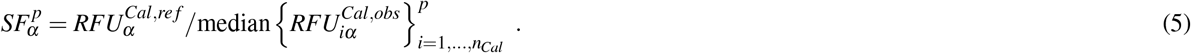

Calibration scale factors may be pinned to an external reference, but here we utilize an internal reference determined by the median across all calibrators on all *n*_*p*_ plates, i.e.

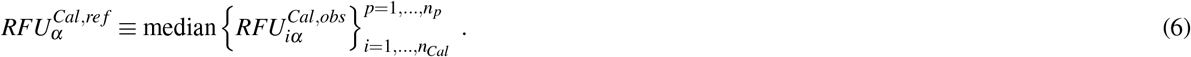

In order to correct the overall brightness level of the *p*-th plate, we calculate the plate-scale scale factor as the median of 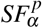 across all SOMAmers, i.e.

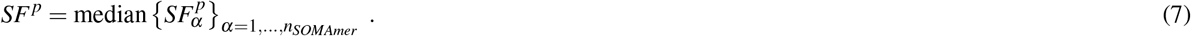

For all wells on the *p*-th plate and all SOMAmers, RFUs are multiplied by the plate-scale factor *SF*^*p*^.

The following code chunk implements the plate-scale normalization step:

~~~
59 RFU[[4]] = RFU[[3]]
60 SF = **matrix**(**rep**(NA,n**_**plate^**\***^n**_**somamer),**ncol**=n**_**somamer)
61 sel**_**cal = sampl[,”SampleType”]==“Calibrator”
62 cal**_**interplate**_**ref = **rep**(NA,n**_**somamer)
63 **for** (i**_**somamer in 1:n**_**somamer) {
64     cal**_**interplate**_**ref[i**_**somamer] = **median**(RFU[[4]][sel**_**cal,i**_**somamer])
65     **for** (i**_**plate in 1:n**_**plate) {
66         sel**_**plate = sampl[,”PlateId”]==plate[i**_**plate]
67         cal**_**intraplate**_**ref = **median**(RFU[[4]][sel**_**cal&sel**_**plate,i**_**somamer])
68         SF[i**_**plate,i**_**somamer] = cal**_**interplate**_**ref[i**_**somamer]**/**cal**_**intraplate**_**ref
69     }
70 }
71 SF**_**plateScale = **apply**(SF,1,**median**)
72 **for** (i**_**plate in 1:n**_**plate) {
73     sel**_**plate = sampl[,”PlateId”]==plate[i**_**plate]
74     RFU[[4]][sel**_**plate,] = RFU[[4]][sel**_**plate,]^**\***^SF**_**plateScale[i**_**plate]
75 }
~~~

### 4. Inter-plate calibration (hyb.msnCal.ps.cal)

Following plate-scale normalization, we recalculate SOMAmer- and plate-specific scale factors via Eqs. (5)-(6). Separately for each SOMAmer and plate, all wells on the plate are corrected by the recalculated 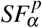.

The following code chunk implements the inter-plate calibration step:

~~~
76 RFU[[5]] = RFU[[4]]
77 SF**_**cal = **matrix**(**rep**(NA,n**_**plate^**\***^n**_**somamer),**ncol**=n**_**somamer)
78 sel**_**cal = sampl[,”SampleType”]==“Calibrator”
79 cal**_**interplate**_**ref**_**N = **rep**(NA,n**_**somamer) *# new interplate reference*
80 **for** (i**_**somamer in 1:n**_**somamer) {
81     cal**_**interplate**_**ref**_**N[i**_**somamer] = **median**(RFU[[5]][sel**_**cal,i**_**somamer])
82     **for** (i**_**plate in 1:n**_**plate) {
83         sel**_**plate = sampl[,”PlateId”]==plate[i**_**plate]
84         cal**_**intraplate**_**ref = **median**(RFU[[5]][sel**_**cal&sel**_**plate,i**_**somamer])
85         SF**_**cal[i**_**plate,i**_**somamer] = cal**_**interplate**_**ref**_**N[i**_**somamer]**/**cal**_**intraplate**_**ref
86         RFU[[5]][sel**_**plate,i**_**somamer]=RFU[[5]][sel**_**plate,i**_**somamer]^**\***^SF**_**cal[i**_**plate,i**_**somamer]
87     }
88 }
~~~

### 5. Median signal normalization on all sample types (hyb.msnCal.ps.cal.msnAll)

At this stage, after correcting plate-to-plate variability to the fullest extent possible, median signal normalization (described in step 2 and Fig. 5) can be performed separately on each sample type. This step yields the final, fully-normalized dataset. The following code chunk implements the median signal normalization step on all sample types:

~~~
89 RFU[[6]] = RFU[[5]]
90 SF**_**medNormFull = **matrix**(**rep**(1,n**_**sampl^**\***^n**_**dil),**ncol**=n**_**dil)
91 sampl**_**type=**c**(“QC”,”Sample”,”Buffer”,”Calibrator”)
92 n**_**sampl**_**type = **length**(sampl**_**type)
93 **for** (i**_**sampl**_**type in 1:n**_**sampl**_**type) {
94      indx**_**sampl = **which**(sampl[,”SampleType”]==sampl**_**type[i**_**sampl**_**type])
95      **for** (i**_**dil in 1:n**_**dil) {
96           sel**_**somamer = **as.numeric**(somamer[,”Dilution”])==dil[i**_**dil]
97           dat = RFU[[6]][indx**_**sampl,sel**_**somamer,**drop**=F]
98           dat2 = dat
99           **for** (i in 1:**ncol**(dat2)) {
100               dat2[,i] = dat2[,i]**/median**(dat2[,i])
101           }
102           **for** (i in 1:**nrow**(dat)) {
103               SF**_**medNormFull[indx**_**sampl[i],i**_**dil] = 1**/median**(dat2[i,])
104               dat[i,] = dat[i,]^**\***^SF**_**medNormFull[indx**_**sampl[i],i**_**dil]
105           }
106           RFU[[6]][indx**_**sampl,sel**_**somamer] = dat
107      }
108 }
~~~

Finally, the following code chunk saves all sample-, SOMAmer-, and plate-based metadata (including the scale factors used in the normalization procedure) along with the RFU matrices at each normalization step:

~~~
109 header=**c**(**colnames**(sampl),”SF**_**hyb”,paste0(“SF**_**msnCal**_**d”,dil**_**lab),paste0(“SF**_**msnAll**_**d”,dil**_**lab))
110 output = **rbind**(header,**cbind**(sampl,SF**_**hyb,SF**_**medNormInt,SF**_**medNormFull))
111 **write**(**t**(output),**ncol**=**ncol**(output),**file**=“samples**_**SF.txt”,sep=“\t”)
112 header=**c**(**colnames**(somamer),”Cal**_**Interplate**_**Ref**_**Pass1”,”Cal**_**Interplate**_**Ref**_**Pass2”)
113 output = **rbind**(header,**cbind**(somamer,cal**_**interplate**_**ref,cal**_**interplate**_**ref**_**N))
114 **write**(**t**(output),**ncol**=**ncol**(output),**file**=“somamers**_**SF.txt”,sep=“\t”)
115 header=**c**(“Plate”,”SF**_**plateScale”,paste0(“SF**_**cal**_**”,somamer[,”SeqId”]))
116 output = **rbind**(header,**cbind**(plate,SF**_**plateScale,SF**_**cal))
117 **write**(**t**(output),**ncol**=**ncol**(output),**file**=“plates**_**SF.txt”,sep=“\t”)
118 **for** (i**_**norm in 2:n**_**norm) {
119     outfile = paste0(“RFU.”,norm[i**_**norm],”.txt”)
120     **write**(**t**(RFU[[i**_**norm]]),**ncol**=n**_**somamer,**file**=outfile,sep=“\t”)
121 }
~~~

It should be pointed out that SomaLogic currently delivers normalized output files (as plain text files of extension `adat`) that follow similar steps as those described here, but using external references^8^. These references may be outdated (because they are specific to the control samples used), not necessarily representative of the target samples of interest (because they utilize a fixed pool of healthy human control samples for the last normalization step) and are not delivered with the `adat` files provided to the customer (therefore precluding any attempt of independent analysis). In contrast, the process described here relies solely on internal references derived from the measured data. In our study of nearly 1,800 experimental samples from the Baltimore Longitudinal Study on Aging (BLSA), we found that fully normalized datasets using internal versus external references were highly concordant. For each human protein SOMAmer in the plasma 7K assay, we calculated the Spearman’s correlation between the fully normalized RFU values from the `adat` file provided by SomaLogic using external references (a file designated with the “hybNorm.medNormInt.plateScale.calibrate.anmlQC.qcCheck.anmlSMP” suffix) and the full normalization described here using internal references (“hyb.msnCal.ps.cal.msnAll”). The distribution of correlation estimates over all 7,289 human protein SOMAmers is shown in Fig. 6; the distribution median is *r* = 0.996. It should be added that, due to Spearman’s correlation invariance, these results remain valid under any type of monotonic (e.g. logarithmic) RFU transformations.

**Figure 6.**
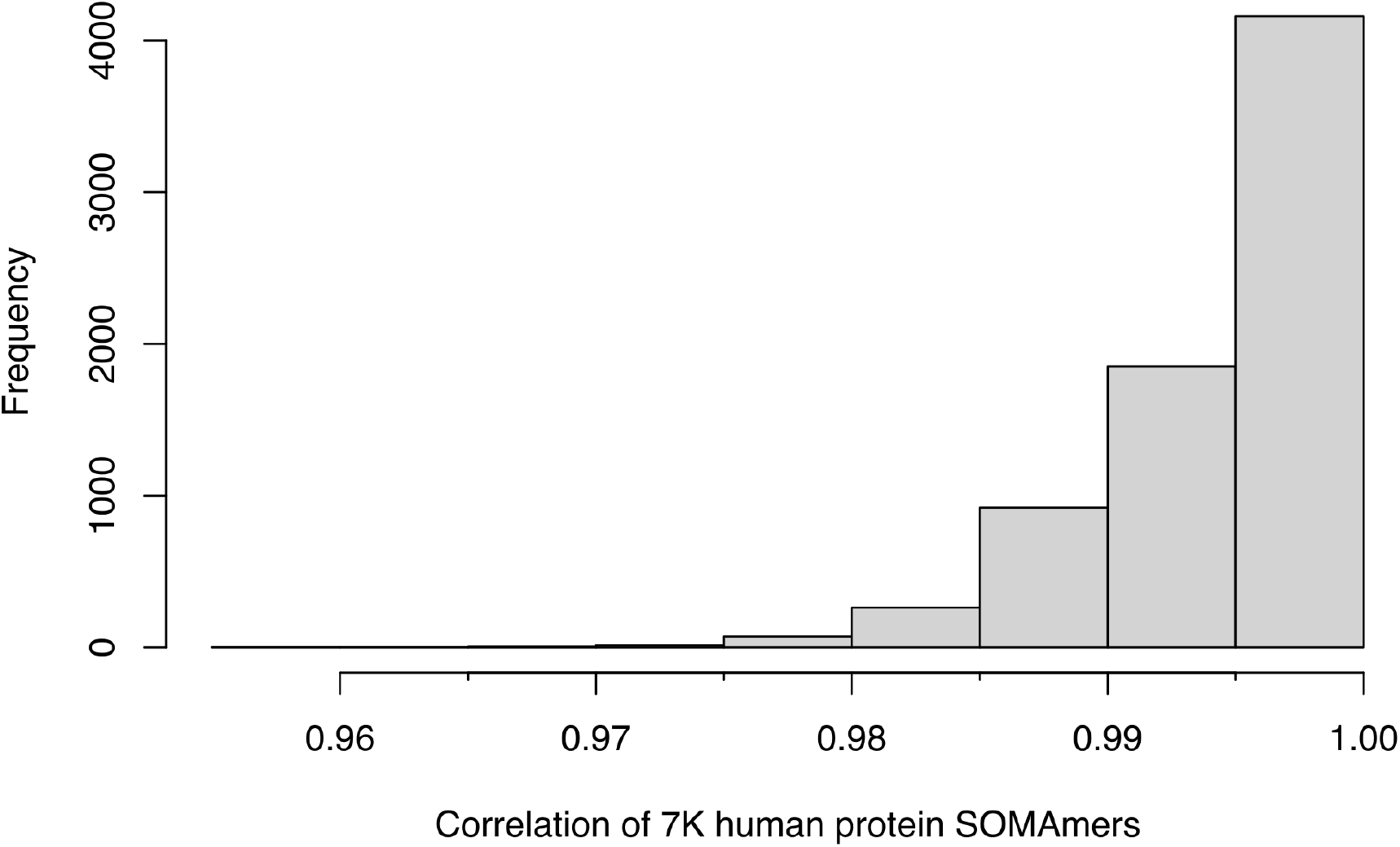
Concordance between normalization approaches. Distribution of Spearman’s correlation estimates for the 7,289 human protein SOMAmers in the 7K plasma SomaScan assay, calculated over nearly 1,800 human donor samples from the BLSA^22^. For each SOMAmer, the correlation was calculated between the fully normalized RFU values from SomaLogic’s pipeline using external references and the full normalization described here using internal references.

SomaLogic’s standard data quality reports utilize scale factors from the normalization procedure to flag samples, SOMAmers, and plates that do not pass pre-established acceptance criteria. Additional standard approaches that SomaScan users may implement are dimensional reduction techniques such as Principal Component Analysis (PCA), which serves both as a quality control and exploration tool to gain insight on the data. Fig. 7 shows PCA performed on 2,050 raw (not normalized) samples, including 68 buffers, 110 calibrators, 66 QC and 2,806 human donor samples from the BLSA^22^. Seven samples, flagged in SomaLogic’s data quality report due to their out-of-range normalization scale factors, are shown by red arrows. In this case, PCA helps to confirm that those samples indeed appear as outliers. Four of them, in fact, appear closer to the reagent-only buffer wells than the large cluster formed by calibrator, QC, and the remaining biological samples. Since this study spans 22 plates, shown by different colors, PCA may provide visual hints of plate effects; inspecting and comparing PCA plots with different normalizations provides more clues to explore whether those potential plate effects were effectively removed by the normalization process^22^. It should be noticed that other bioinformatic tools, developed to assess and remove batch effects for other omics, may be useful in the context of SomaScan data processing, for instance: ComBat^26^, implemented in the sva R package^27^, a very well established method for batch correction in microarray and RNA-seq data; guided PCA^28^; and multi-MA normalization^29^, among others. This important topic certainly deserves further investigation.

**Figure 7.**
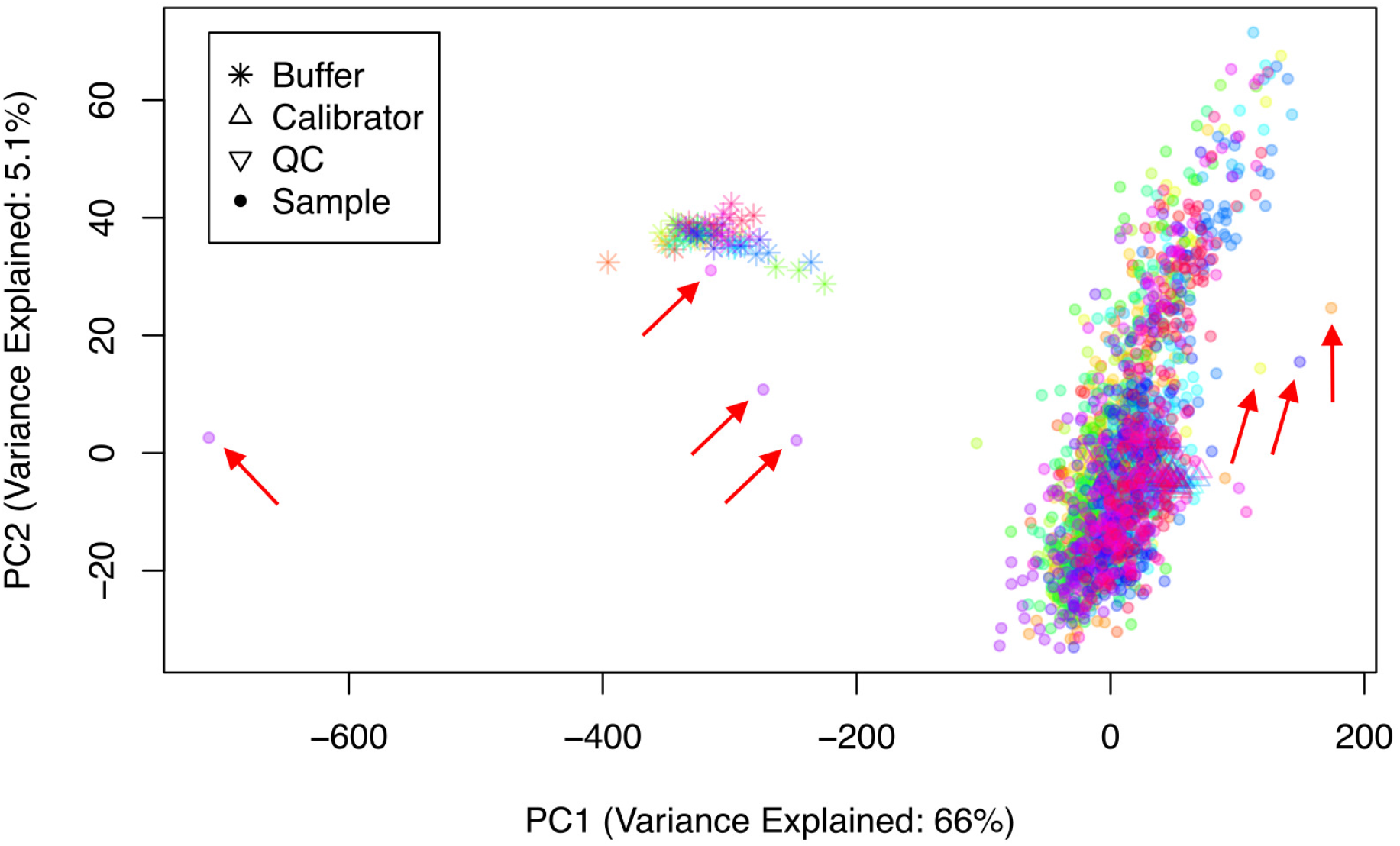
PCA as quality control and data exploration tool. PCA performed on 2,050 raw (not normalized) samples, including 68 buffers, 110 calibrators, 66 QC and 2,806 BLSA samples distributed across 22 plates. Each plate is shown by a different color. The red arrows show seven samples flagged due to their out-of-range normalization scale factors.

## Pre-Analytical Variation (PAV) SomaSignal Tests (SSTs)

Pre-analytical variation (PAV) due to sample collection, handling, and storage is known to affect many analyses in molecular biology. By implementing data modeling techniques similar to those previously developed to find SomaScan signatures associated with clinical phenotypes^8^, SomaLogic has developed a novel set of so-called SomaSignal Tests (SSTs) to assess pre-analytical variation due to different sample processing factors, including fed-fasted time, number of freeze-thaw cycles, time-to-decant, time-to-spin, and time-to-freeze. While it is not always possible to control sample collection, processing, storage, and handling to ensure consistency, PAV-SSTs were designed to quantitatively estimate whether and to what extent samples may have been impacted by these nuisance factors.

In previous work^22^, we implemented a framework to assess technical variability based on measurements performed on duplicate biological inter-plate pairs from samples obtained from *n*_*dupl*_ = 102 human participants in the BLSA. Panels (a)-(e) of Fig. 8 show, for each PAV metric, the magnitude of the difference between estimates from each duplicate pair, Δ*x*_*i*_ = |*x*_*i*,1_*− x*_*i*,2_ |, as a function of their average,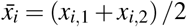. Next, we will leverage this framework to generate an independent assessment of the robustness of SomaLogic’s PAV estimates. Let us first recall that, based on the scaled relative differences, defined as 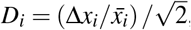, different variability estimates may be built, namely: (i) the Root-Mean-Squared Variation (RMSV) metric, defined as

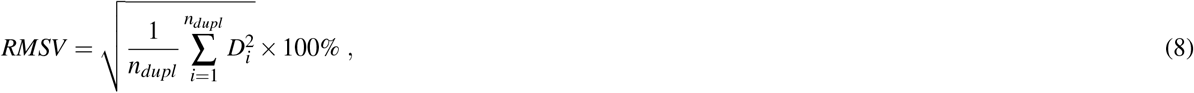

(ii) the Mean Absolute Difference Variation (MADV) metric, defined as

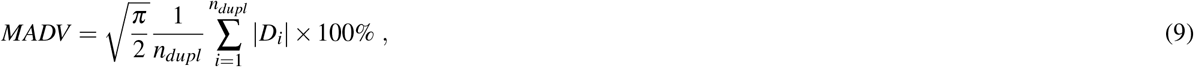

and (iii) the Percentile Variation (PV) metric, defined as

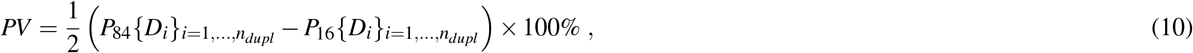

where *P*_84_ and *P*_16_ are the 84^th^ and 16^th^ percentiles in the distribution of scaled relative differences. If the scaled relative differences are normally distributed, these variability estimates are equivalent; the magnitude of the differences between these estimates is indicative of the magnitude and severity of deviations from normality. Armed with these various definitions, Panel Fig. 8(f) shows the percent variation across PAV estimates. As expected by theoretical considerations, the PV metric is the least sensitive to distribution tails and therefore yields the lowest variation. We observe that, with the exception of the time-to-freeze estimate, which shows a PV variation of about 15%, the PV variation for the remaining four PAV metrics lies approximately within the 5 *−* 7% range.

**Figure 8.**
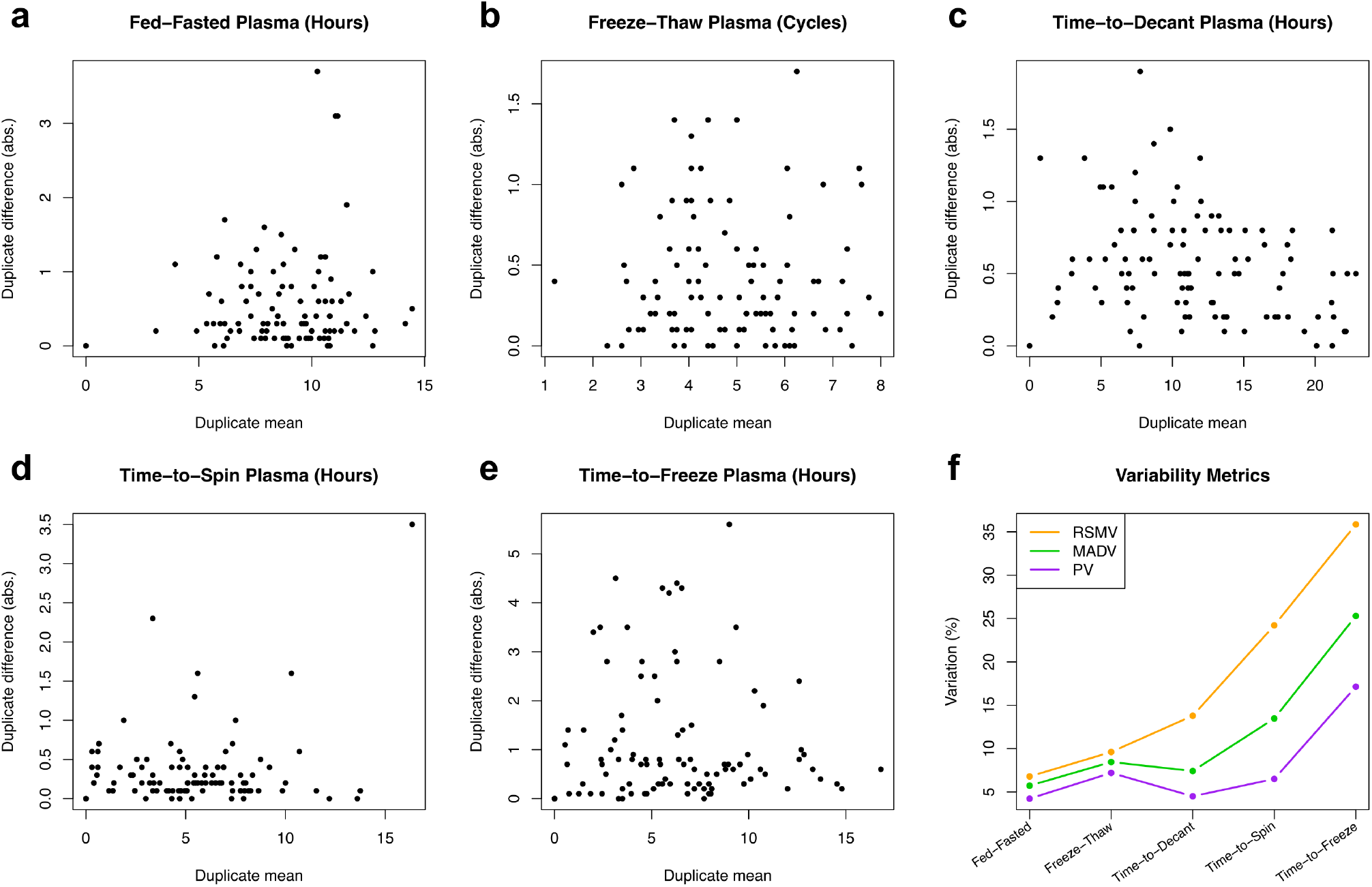
Assessment of the robustness of PAV-SST estimates using BLSA technical replicates. **a-e** Duplicate difference vs duplicate mean for different PAV-SST estimates, as indicated. Each circle represents one pair of technical duplicates from the same human donor sample. **f** Variation of PAV-SST estimates determined by different metrics, namely: Root-Mean-Squared Variation (RMSV), Mean Absolute Difference Variation (MADV), and Percentile Variation (PV).

## Summary and Conclusions

With the addition of the latest v5.0 (11K) version released on November 1, 2023, SomaScan galvanized its role as one of the leading technologies in high-throughput human proteomics. Going back to the v3 (1.3K) version about eight years prior, the assay has shown a steady, approximately linear growth in protein coverage. Several independent teams, including ours at the NIH, have explored the assay’s sensitivity and variability^16–22^. Furthermore, multiple studies have investigated SomaScan’s specificity, cross-reactivity, and orthogonal assay reproducibility^4,5,30–32^. Less attention, however, has been dedicated to explore normalization procedures. Typically, SomaLogic provides one or several data files in `adat` format, which are normalized using their internal analysis pipeline. This pipeline, however, has historically remained under SomaLogic’s control and operated to a large extent as a black box, providing SomaScan users only with descriptive explanations of the steps involved in the process. SomaLogic’s normalization procedures, moreover, have evolved overtime, switching from internal references to external references, altering the order in which median normalization was implemented, and splitting the interplate calibration step into plate-level rescaling (referred to as plate-scale normalization) followed by SOMAmer- and plate-specific normalization. To the best of our knowledge, no software version control of these changes have been shared with SomaScan users, making it difficult the task to compare or integrate studies run years apart, perhaps even using different versions of the assay.

As an early contribution to the nascent field of SomaScan bioinformatics, our NIH labs have undertaken the task to reconstruct the steps involved in a normalization procedure that, without the use of SomaLogic’s proprietary external references, is capable of reproducing normalized RFU values concordant with those from SomaLogic’s pipeline^16,22^. The latest version of our normalization pipeline is described here in detail, including its R code implementation, and available from a public repository. The ability to run independent normalization analyses is important for various reasons. On the one hand, the set of external references (based on a fixed pool of healthy human control samples) may be inappropriate to study individuals and populations with large deviations from a healthy plasma proteome. As noted by Lopez-Silva et al^33^ in their study of chronic kidney disease, SomaLogic’s normalization could potentially attenuate the strength of associations by reducing more extreme values if they are associated with a clinical outcome, perhaps contributing to the attenuated prognostic associations they observed with SomaScan. Pietzner et al^20^ report that using SomaScan data without a normalization step applied to correct for unwanted technical variation and to make data comparable across cohorts, a higher median correlation with results from Olink was observed, along with substantial differences in the association with various phenotypic characteristics. On the other hand, users may be interested in using their own set of control samples to bridge across studies, thereby needing to calibrate different studies using those controls. More generally, users may be interested in adapting the normalization process according to their studies’ characteristics and objectives (for instance, splitting the median normalization procedure into different sample subclasses identified by clinical phenotype). Therefore, we believe that presenting this independent normalization pipeline empowers SomaScan end users to tailor the normalization procedures to their own individual needs.

Following our discussion of normalization approaches, we briefly described quality control procedures based on combining flags from outlier normalization scale factors (such as those provided in SomaLogic’s standard data quality reports) with Principal Component Analysis (PCA), a linear dimensional reduction technique. Alternatively, non-linear dimensional reduction methods such as Uniform Manifold Approximation and Projection (UMAP) and t-distributed Stochastic Neighbor Embedding (t-SNE) could be used for the purpose of quality control and data exploration. In addition, we suggested that bioinformatic tools developed to assess and remove batch effects in data from other omics, such as ComBat^26^, Surrogate Variable Analysis (SVA)^27^, guided PCA^28^, and multi-MA normalization^29^, may be useful techniques that deserve further investigation.

Finally, we investigated SomaLogic’s SomaSignal Test (SST) models, which were recently developed to estimate a variety of Pre-Analytical Variation (PAV) metrics, including fed-fasted time, number of freeze-thaw cycles, time-to-decant, time-to-spin, and time-to-freeze. Leveraging technical duplicates from our study of human donors from the Baltimore Longitudinal Study on Aging (BLSA), we performed an independent evaluation of the statistical variation of the PAV estimates. Within the limited scope of our assessment, we conclude that PAV-SST estimates appear to be robust (percentile variation in the 5*−*7% range, with the exception of time-to-freeze, which is about 15%) and may be used during biomarker evaluations to exclude samples due to nuisance effects related to sample collection and processing, and/or to incorporate these estimates as model covariates to control the downstream impact of those unwanted effects.

## Data and Software Availability

Anonymized datasets and R source code for the normalization pipeline described here are available on the Open Science Framework repository, osf.io/srgef. DOI 10.17605/OSF.IO/SRGEF.

## Acknowledgements

This work was supported entirely by the Intramural Research Program of the National Institute on Aging (NIA).

